# A gene based approach to test genetic association based on an Optimally Weighted Combination of Multiple Traits

**DOI:** 10.1101/592568

**Authors:** Jianjun Zhang, Qiuying Sha, Guanfu Liu, Xuexia Wang

## Abstract

There is increasing evidence showing that pleiotropy is a widespread phenomenon in complex diseases for which multiple correlated traits are often measured. Joint analysis of multiple traits could increase statistical power by aggregating multiple weak effects. Existing methods for multiple trait association tests usually study each of the multiple traits separately and then combine the univariate test statistics or combine p-values of the univariate tests for identifying disease associated genetic variants. However, ignoring correlation between phenotypes may cause power loss. Additionally, the genetic variants in one gene (including common and rare variants) are often viewed as a whole that affects the underlying disease since the basic functional unit of inheritance is a gene rather than a genetic variant. Thus, results from gene level association test can be more readily integrated with downstream functional and pathogenic investigation, whereas many existing methods for multiple trait association tests only focus on testing a single common variant rather than a gene. In this article, we propose a statistical method by Testing an Optimally Weighted Combination of Multiple traits (TOW-CM) to test the association between multiple traits and multiple variants in a genomic region (a gene or pathway). We investigate the performance of the proposed method through extensive simulation studies. Our simulation studies show that the proposed method has correct type I error rates and is either the most powerful test or comparable with the most powerful tests. In addition, we illustrate the usefulness of TOW-CM by analyzing a whole-genome genotyping data from a COPDGene study.

## Introduction

Complex diseases are often characterized by many correlated phenotypes which can better reflect their underlying mechanism. For example, hypertension can be characterized by systolic and diastolic blood pressure (Newton-Cheh et al. 2009); metabolic syndrome is evaluated by four component traits: high-density lipoprotein (HDL) cholesterol, plasma glucose and Type 2 diabetes, abdominal obesity, and diastolic blood pressure (Zabaneh and Balding 2010); and a person’s cognitive ability is usually measured by tests in domains including memory, intelligence, language, executive function, and visual-spatial function (Yang and Wang 2012). Also, more and more large cohort studies have collected or are collecting a broad array of correlated phenotypes to reveal the genetic components of many complex human diseases. Therefore, by jointly analyzing these correlated traits, we can not only gain more power by aggregating multiple weak effects, but can also understand the genetic architecture of the disease of interest (Solovieff et al., 2013).

Even though genome-wide association studies (GWASs) have been remarkably successful in identifying genetic variants associated with complex traits and diseases, the majority of the identified genetic variants only explain a small fraction of total heritability (Manolio et al., 2009). On the other hand, a gene is the basic functional unit of inheritance whereas the GWAS are primarily focused on the paradigm of single common variant. However, most published GWAS only analyzed each individual phenotype separately, although results on related phenotypes may be reported together. Large-scale GWAS of complex traits have consistently demonstrated that, with few exceptions, common variants have moderate-to-small effects. Thus, it is important to identify appropriate methods that fully utilize information in multivariate phenotypes to detect novel genes in genetic association studies.

In GWAS, several methods have been developed for multivariate phenotypes association analysis (Yang and Wang 2012) to test association between multivariate continuous phenotypes and a single common variant. To our knowledge, current multivariate phenotypes association methods can be roughly classified into two categories: univariate analysis and multivariate analysis. Univariate analysis methods perform an association test for each trait individually and then combines the univariate test statistics or combine the p-values of the univariate tests (Yang et al. 2010, O’Brien 1984, Kim et al. 2015 and Liang et al. 2016). Even though this method is very convenient to compute and analysis, it ignores the potential correlation among all individual phenotypes and may reduce the power compared to multivariate analysis. Multivariate analysis methods jointly analyze more than one phenotype in a unified framework and test for the association between multiple phenotypes and genetic variants. Multivariate analysis methods include multivariate analysis of variance (MANOVA) (Cole et al. 1994), linear mixed effect models (LMM) (Laird and Ware 1982), and generalized estimating equations (GEE) (Liang and Zeger 1986) method. Another special approach is to consider reducing the dimension of the multivariate phenotypes by using dimension reduction techniques. The common method for dimensionality reduction is principal component analysis (PCA) (Ott and Rabinowitz, 1999) which essentially finds the combination of these phenotypes and assumes that the transformed phenotypes are independent. The limitation of this method is that it can not properly account for the variation of phenotypes or genotypes, and also it is hard to interpret the meaning of principle components of the multivariate phenotypes, especially in practice.

Recent studies show that complex diseases are caused by both common and rare variants (Pritchard 2001; Pritchard and Cox 2002; Walsh and King 2007; Bodmer and Bonilla 2008; Stratton and Rahman 2008; Kang et al. 2010; Teer and Mullikin 2010). Gene-based analysis requires statistical methods that are fundamentally different from association statistics used for testing common variants. It is essential to develop novel statistical method to test the association between multiple traits and multiple variants (common and/or rare variants). In this article, we develop a statistical method to test the association between multiple traits and genetic variants (rare and/or common) in a genomic region by Testing the association between an Optimally Weighted combination of Multiple traits (TOW-CM)and the genomic region. TOW-CM is based on the score test under a linear model, in which the weighted combination of phenotypes is obtained by maximizing the score test statistic over weights. The weights at which the score test statistic reaches its maximum are called the optimal weights. We also use extensive simulation studies to compare the performance of TOW-CM with MANOVA (Cole et al. 1994), MSKAT (Wu and Pankow 2016) and minimum p-value (Sha et al., 2012). Simulation studies demonstrate that, in all the simulation scenarios, TOW-CM is either the most powerful test or comparable to the most powerful test among the four tests. We also illustrate the usefulness of TOW-CM by analyzing a real whole-genome genotyping data from a COPDGene study.

## Methods

We consider a sample with *n* unrelated individuals. Each individual has *K* (potentially correlated) traits and has been genotyped at *M* variants in a considered region (a gene or a pathway). Denote *y*_*ik*_ as the *k*^*th*^ trait value of the *i*^*th*^ individual and *x*_*im*_ as the genotype score in additive coding of the *i*^*th*^ individual at the *m*^*th*^ variant. Let ***Y*** = (*Y*_1_, · · ·, *Y*_*K*_) denote the random vector of *K* traits and ***X*** = (*X*_1_, · · ·, *X*_*M*_) denote the random variable of the genotype score at *M* variants for these *n* individuals where *Y*_*k*_ = (*y*_1*k*_, · · ·, *y*_*nk*_)^*T*^ and *X*_*m*_ = (*x*_1*m*_, · · ·, *x*_*nm*_)^*T*^. Consider a linear combination of ***Y*** denoted as 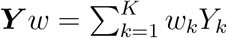, where *w* = (*w*_1_, · · ·, *w*_*K*_)^*T*^.

We model the relationship between the combination of multiple continuous traits with the *M* genetic variants in the considered region using linear model

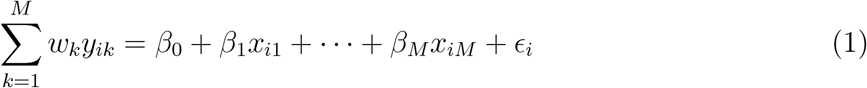

where *β*_0_ is the intercept and ***β*** = (*β*_1_, · · ·, *β*_*M*_)^*T*^ is the corresponding vector of coefficient. To test the association between the combination of the multiple traits with the *M* genetic variants is equivalent to test the null hypothesis *H*_0_: ***β*** = 0 under equation (1). We use the score test statistic given by Sha et al., (2011) to test *H*_0_: ***β*** = 0 under equation (1). Let 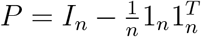 and the test statistic is:

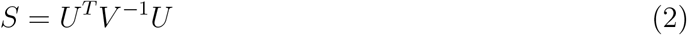

where *U* = (*P **X***)′ *P **Y** w* and 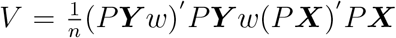. The score test can be rewritten as a function of *w*:

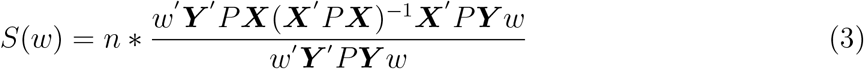

where *P* = *P*′ and *PP*′ = *P*. We propose to maximize *S*(*w*) to get the optimal weight and then define the statistic to test the effect between the optimally weighted combination of traits with genetic variants.

When *D* = ***Y**′ P**Y*** is positive definite, maximizing *S*(***w***) is equivalent to maximizing

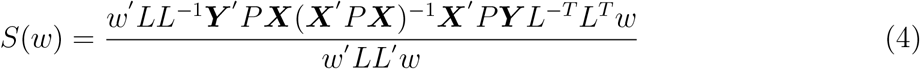

where *L* is the lower triangular matrix obtained from the Cholesky decomposition of *D* = *LL*^*T*^. However, the matrix of *D* is usually not full rank because of existing correlation between multiple traits. If the matrix *D* is semi-positive define matrix, we introduce a ridge parameter λ_0_, for which we suggest the choice 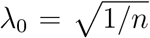, where *n* is the number of individuals in the testing data, and modify the adjustment to mitigate the effect of the non-positive matrix *D* in order to avoid the instability: *D* = ***Y***′ *P**Y*** + λ_0_*I*. Let *C* = *L*^−1^***Y*′** *P**X***(***X*′** *P **X***)^−1^ ***X*′***P**Y** L*^−*T*^ and ***c*** be the eigenvector corresponding to the largest eigenvalue of the matrix *C*, then *S*(***w***) is maximized when *L′* ***w*** equals to ***c***. Hence equation (4) is maximized when *w*^*o*^ = *L*^−*T*^ ***c***. Specially, if all the traits we consider are independent and *M* = 1, we can get an analytical weight referred to (Sha et al., 2012):

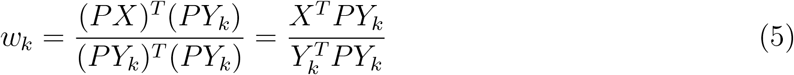

for the *k*^*th*^ phenotype, *k* = 1, 2, 3, …, *K*. The equation (5) is equivalent to 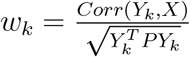 where the numerator is the correlation coefficient between the *k*^*th*^ phenotype *Y*_*k*_ and the genotypic variant *X* and the denominator can be viewed as the variance of the *k*^*th*^ phenotype *Y*_*k*_. It means that *w*_*k*_ has same direction with the correlation between the phenotype *Y*_*k*_ and the genotypic variant *X*, and puts big weight to the *k*^*th*^ trait when it has strong association with the genotypic variant and/or it has low variance.

We define the statistic to test an optimally weighted combination of multiple traits (TOW-CM), 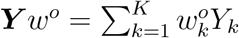 as

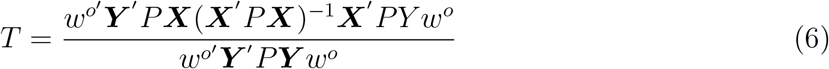

We use permutation methods to evaluate P-values of *T*. The TOW-CM method can also be extended to incorporate covariates. Suppose that there are *p* covariates. Let *z*_*il*_ denote *l*^*th*^ covariate of the *i*^*th*^ individual. We adjust both trait value *y*_*ik*_ and genotypic score *x*_*im*_ for the covariates by applying linear regressions. That is,

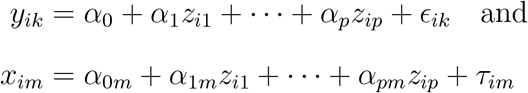

Let 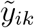 and 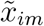 denote the residuals of *y*_*ik*_ and *x*_*im*_, respectively. We incorporate the covariate effects in TOW-CM by replacing *y*_*ik*_ and *x*_*im*_ in equation (6) by 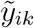 and 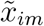. With covariates, the statistic of TOW-CM is defined as:

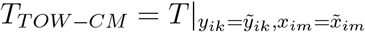

## Comparison of tests

We compared the performance of our method (TOW-CM) with the following methods: 1) Multivariate Analysis of Variance (MANOVA) (Cole et al. 1994); 2) Multi-trait Sequence Kernel Association Test (MSKAT) (Wu and Pankow 2016); 3) Minimum p-value based on the p-values of the individual trait TOW (Sha et al., 2012) significance p-values (denoted as minP).

## Simulation

In simulation studies, we use the empirical Mini-Exome genotype data including genotypes of 697 unrelated individuals on 3205 genes obtained from Genetic Analysis Workshop 17 (GAW17). Two differen type of variants (Common variants: minor allele frequency (MAF)>0.05 and Rare variants: MAF<0.05) are chosen from a super gene (Sgene) including four genes: ELAVL4 (gene1), MSH4 (gene2), PDE4B (gene3), and ADAMTS4 (gene4). The pattern of the allele frequency distribution of Sgene are similar as the 3205 genes (Sha et al., 2012). In our simulation studies, we generate genotypes based on the genotypes of 697 individuals in these four genes. The genotypes are extracted from the sequence alignment files provided by the 1,000 Genomes Project for their pilot3 study (http://www.1000genomes.org). To generate the genotype of an individual, we generate two haplotypes according to the haplotype frequencies.

We test K=4 related traits with a compound-symmetry correlation matrix and consider two covariates: a standard normal covariate *z*_1_ and a binary covariate *z*_2_ with *P* (*z*_2_ = 1) = 0.5. We generate trait values based on genotypes by using the following models:

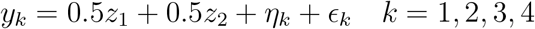

where ***ϵ*** = (*ϵ*_1_, *ϵ*_2_, *ϵ*_3_, *ϵ*_4_) are zero-mean normal with variances 1 and correlation *ρ*. We set the magnitude of correlation |*ρ*| = 0.2, 0.5 and 0.8 and the signs of symmetric location of covariate matrix are randomly chosen from (−1,1). ***η*** = (*η*_1_, *η*_2_, *η*_3_, *η*_4_) are contributions from a set of genotypic variants, which are simulated as follows.

For type I error, phenotypes are generated under the null model i.e. ***η*** = 0. To evaluate power, we randomly choose one common variant and *n*_*c*_ (20%) rare variants as casual variants. We assume that all the *n*_*c*_ rare causal variants have the same heritability and the heritability of the common causal variant is twice of the heritability of rare causal variants. That is, we model the genotypic variants’ contribution to disease risk as 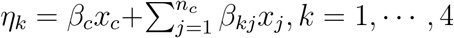 where *x*_*c*_ and *x*_*j*_ denote the common variant and rare variant, respectively. *β*_*c*_ and *β*_*kj*_ represent the corresponding effect size. Let *h* and *h*_*k*_ denote the heritability of all the causal variants for all the*K* traits and for the *k*^*th*^ trait, respectively. We generate *K* random numbers *t*_1_, · · ·, *t*_*K*_ from a uniform distribution between 0 and 1, and the heritability of *k*^*th*^ trait denotes 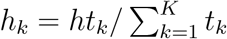. For *k*^*th*^ trait, we assign that the effect size of common variate

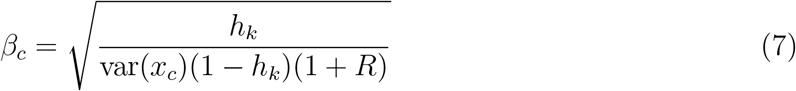

and the magnitude of the effect of rare variate

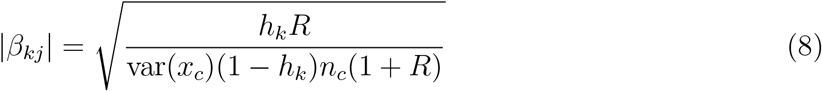

where *R* denote the ratio of the heritability of rare causal variants to the heritability of the common causal variant. For power comparisons, We conduct simulations for four scenarios: each time only the first *L* traits are associated with the rare variant set, *L* = 1, 2, 3, 4, respectively. Intuitively, in the first scenario (L = 1), when only the first trait is associated with the variants set, the minP method (it equals to test the first trait alone) may have good performance. However, we will show that by simultaneously testing correlated null traits, our proposed method (TOW-CM) could actually improve the detection power compared to test the first trait alone. When there are multiple correlated traits that are associated with the rare variants set, the proposed TOW-CM could offer much improved detection power than the minimum p-value based approach. In each scenario, we also consider different percentage of risk variants for rare variants.

## Simulation Results

Table (1) summarizes the estimated type I errors of our method TOW-CM under different significance levels and different magnitude of trait correlation |*ρ*|. The type I error rates are evaluated using 10000 replicated samples and the P-values are estimated using 10000 permutations. For the 10000 replicated samples, the 95% confidence intervals (CIs) for the estimated type I error rates of nominal levels 0.05, 0.01, and 0.001 are (0.046, 0.054), (0.008, 0.012), and (0.0004, 0.0016), respectively. From this table, we can see that all of the estimated type I error rates are either within 95% CIs or close to the bound of the corresponding 95% CIs, which indicate that the type I error rates of the proposed method are controlled under all considered scenarios.

**Table 1:**
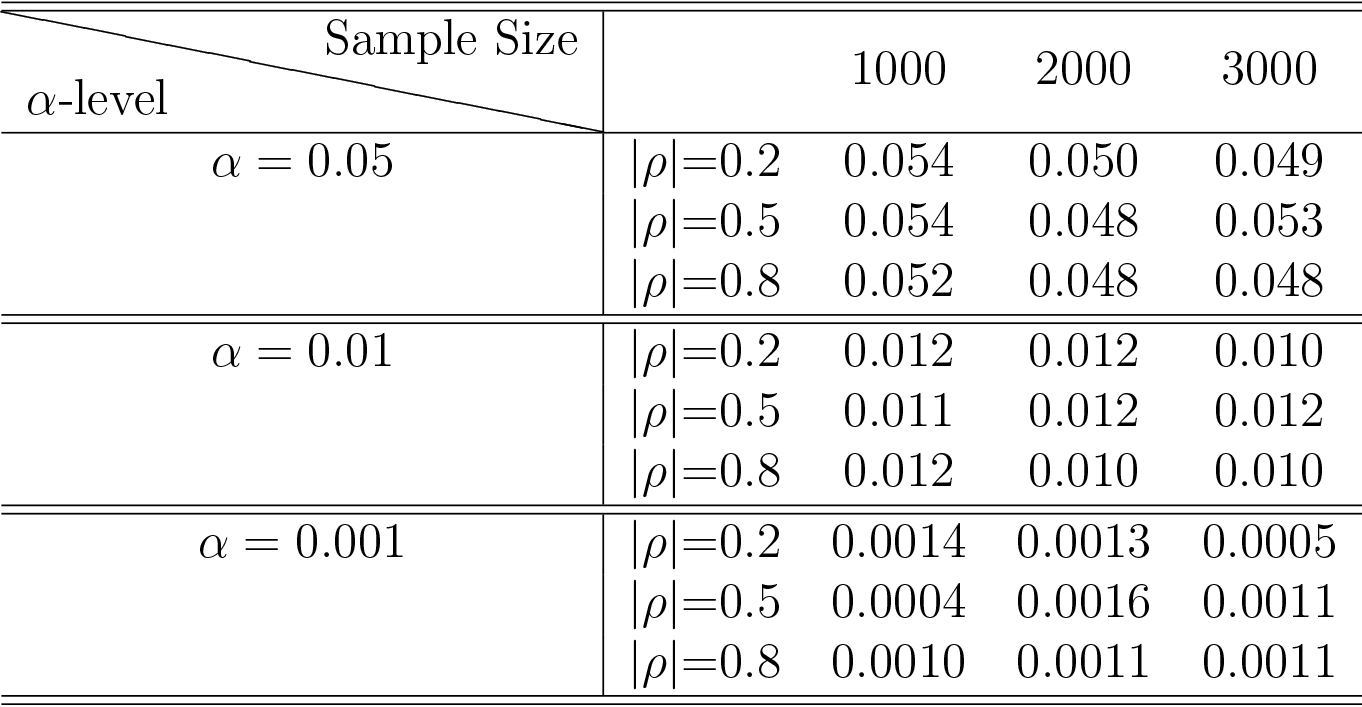
The estimated type I error rates of TOW-CM for four traits under different scenarios

In power comparisons, the P-values of TOW-CM, minP are calculated using 1000 permutations, while the P-values of MANOVA and MSKAT are calculated by asymptotic distributions. The powers of all the four tests are evaluated using 1000 replicated samples at a significance level of 0.05 where Figure 1-6 summarize the power under significance level 0.05 for *L* = 4, 3, 2, 1 respectively.

**Figure 1:**
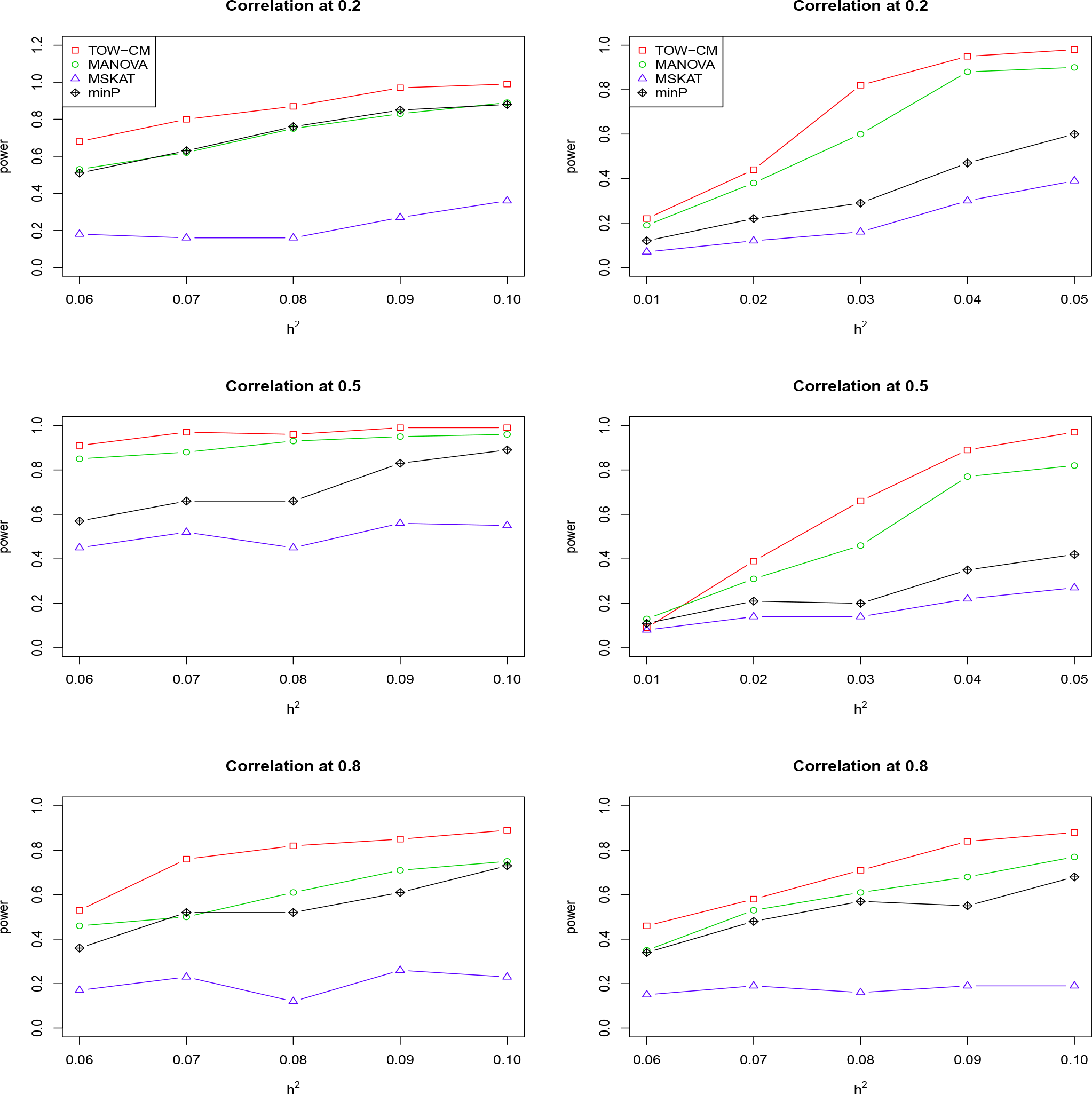
Power Comparison of four tests as a function of heritability for four continuous traits with the magnitude of correlation at 0.2, 0.5 and 0.8, respectively. All the four traits are associated with the gene for the left panel and only first three traits are associated with the gene for the right panel. Sample size is 1,000 and 20% of rare variants are causal. All causal variants are risk variants. The powers are evaluated at a significance level of 0.05.

**Figure 2:**
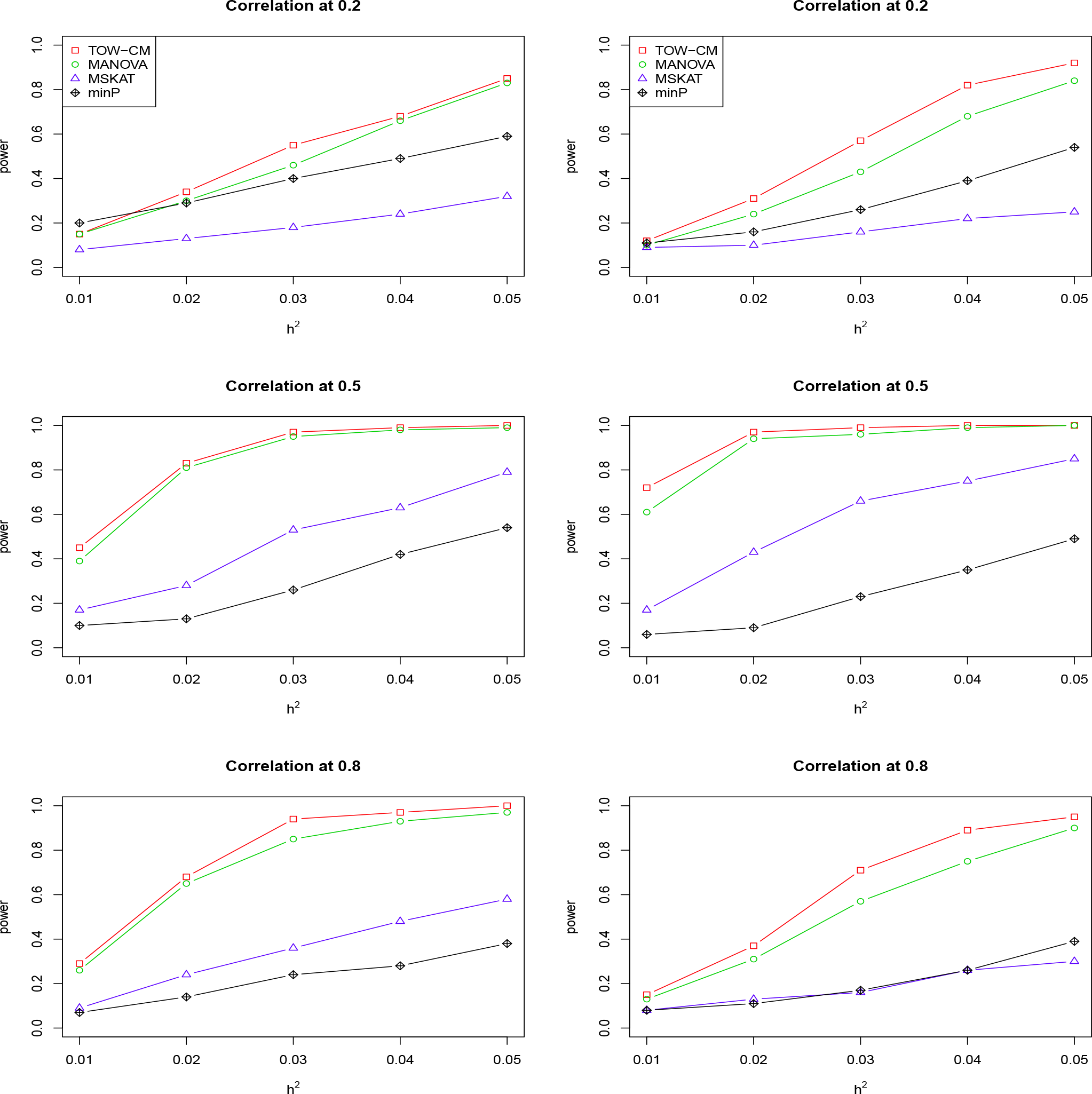
Power comparison of four tests as a function of heritability for four continuous traits with the magnitude of correlation at 0.2, 0.5 and 0.8, respectively. All the four traits are associated with the gene for the left panel and only first three traits are associated with the gene for the right panel. Sample size is 1,000 and 20% of rare variants are causal variants among which 90% of causal variants are risk variants and 10% of causal variants are protective variants. The powers are evaluated at a significance level of 0.05.

**Figure 3:**
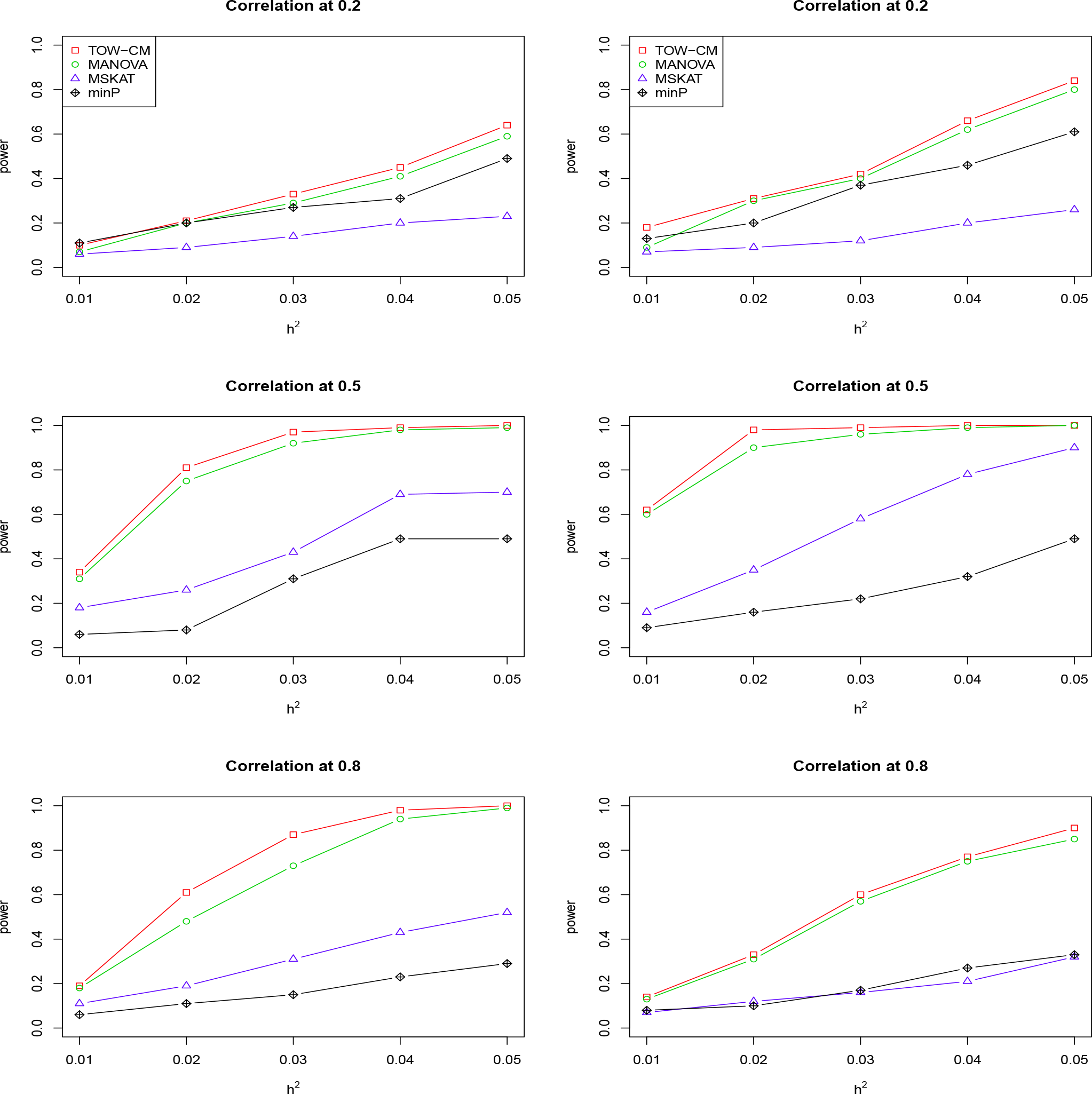
Power comparison of four tests as a function of heritability for four continuous traits with the magnitude of correlation at 0.2, 0.5 and 0.8, respectively. All the four traits are associated with the gene for the left panel and only first three traits are associated with the gene for the right panel. Sample size is 1,000 and 20% of rare variants are causal among which 80% of causal variants are risk variants and 20% of causal variants are protective variants. The powers are evaluated at a significance level of 0.05.

**Figure 4:**
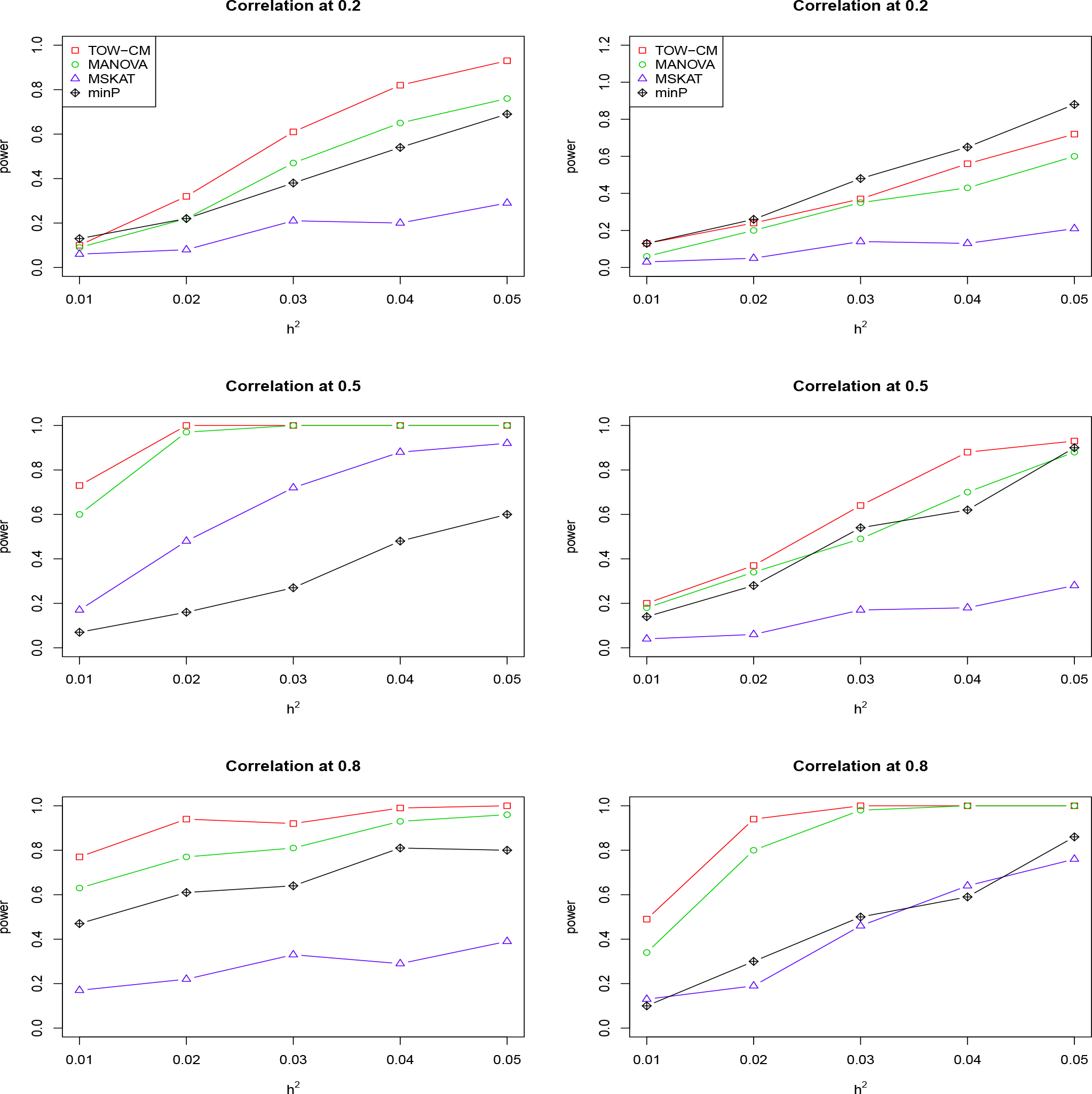
Power comparison of four tests as a function of heritability for four continuous traits with the magnitude of correlation at 0.2, 0.5 and 0.8, respectively. Only first two traits are associated with the gene for left panel and only the first traits are associated with the gene for right panel. Sample size is 1,000 and 20% of rare variants are causal variants. All causal are risk variants. The powers are evaluated at a significance level of 0.05.

**Figure 5:**
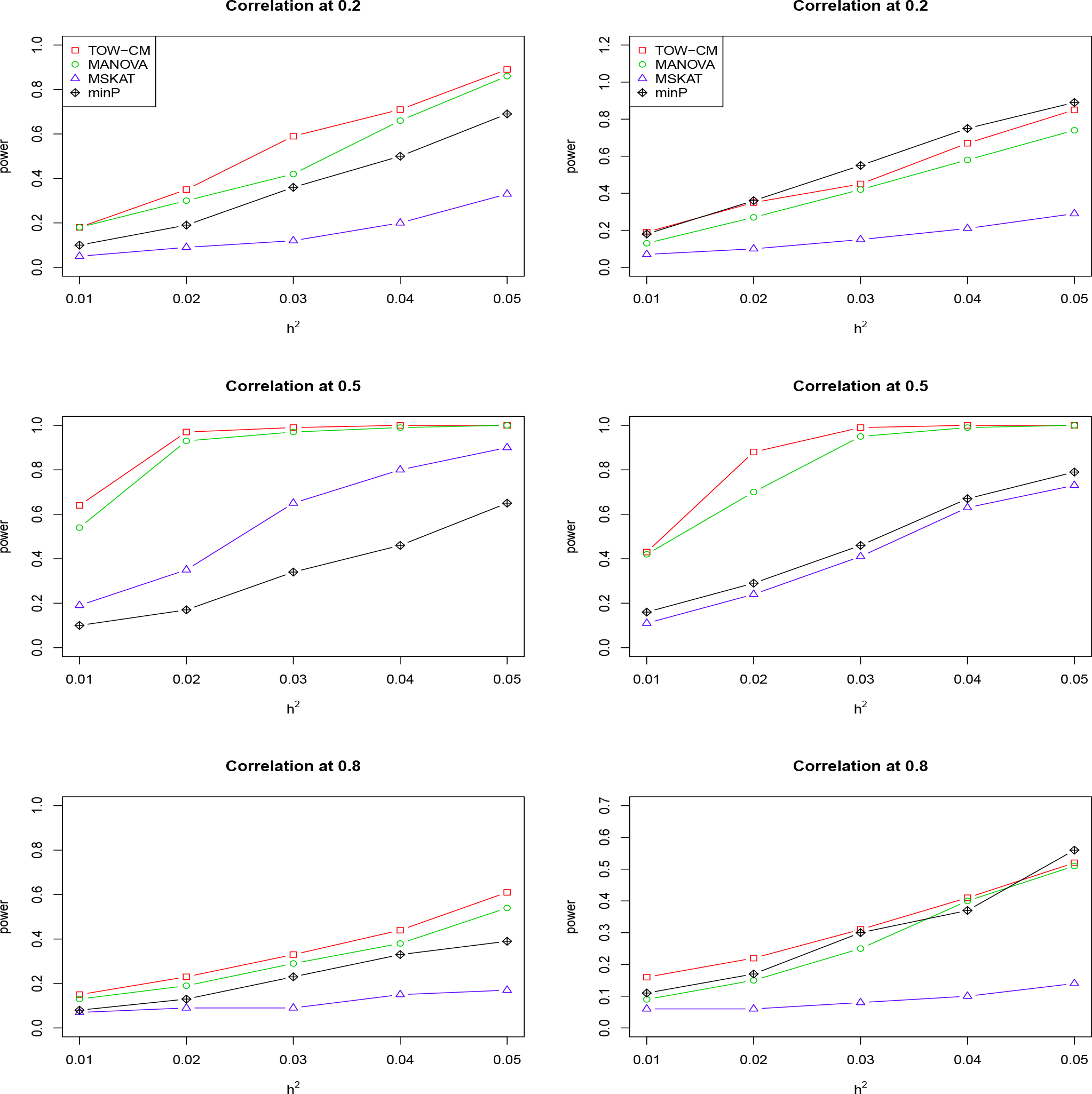
Power comparison of four tests as a function of heritability for four continuous traits with the magnitude of correlation at 0.2, 0.5 and 0.8, respectively. Only first two traits are associated with the gene for left panel and only the first traits are associated with the gene for right panel. Sample size is 1,000 and 20% of rare variants are causal. 90% of causal are risk variants and 10% of causal are protective variants. The powers are evaluated at a significance level of 0.05.

**Figure 6:**
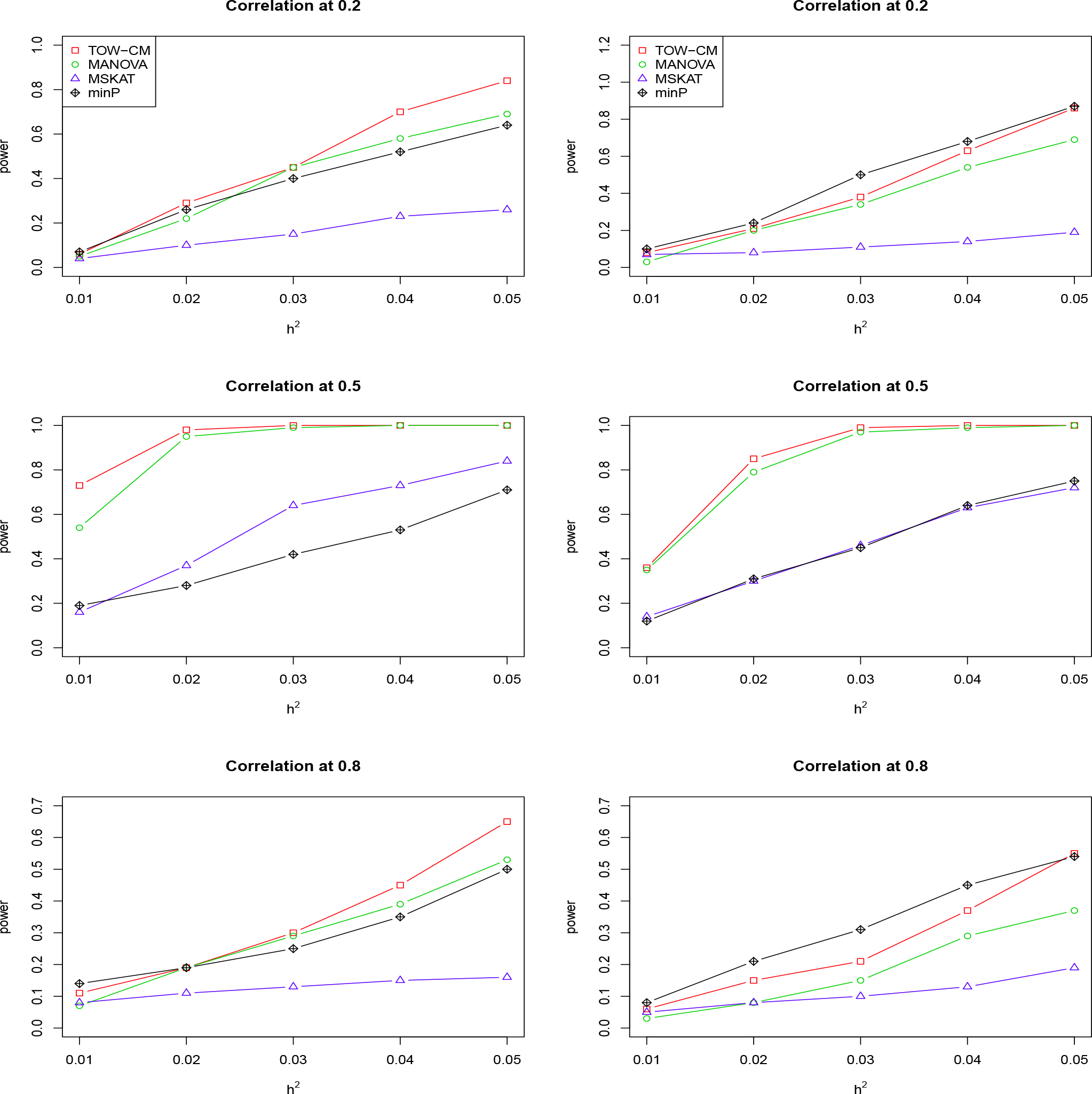
Power comparison of four tests as a function of heritability for four continuous traits with the magnitude of correlation at 0.2, 0.5 and 0.8, respectively. Only first two traits are associated with the gene for left panel and only the first traits are associated with the gene for right panel. Sample size is 1,000 and 20% of rare variants are causal among which 80% of causal variants are risk variants and 20% of causal variants are protective variants. The powers are evaluated at a significance level of 0.05.

These figures show the power comparisons of the four tests (TOW-CM, MANOVA, MSKAT and minP). Power is a function of the total heritability based on three cases (all causal are risk variants, 90% causal are risk variants, and 80% causal are risk variants) for each specific scenario L. These figures show that TOW-CM is consistently the most powerful test among the four tests, and MANOVA is the second most powerful test when genotypes of genetic variants have impact on more than 1 traits. MSKAT is consistently less powerful than the other two multivariate tests (TOW-CM and MANOVA) probably because there are only 8% variants with MAF in the range (0.01,0.035) in Sgene which the simulations are based on. Similar to SKAT, MSKAT will lose power when the MAF of causal variants are not in the range (0.01,0.035) (Yang et al. 2017). The minP method is consistently less powerful than TOW-CM and MANOVA because they ignore the traits’ dependence by directly using minimum of the P-values of testing the four single traits. Overall, we can see that they suffer power loss with increasing traits correlations.

An interesting scenario is one in which only the first trait is associated with the variants set and all the others are null traits (L = 1). Stephens (2013) and Wu and Pankow (2015) have reported that joint testing by incorporating a correlated null trait could improve the power for testing association of a common variant. When only the first trait is associated with the variants set, minP is either the most powerful test or has similar power to the most powerful test especially in the case of both causal variants under weak traits correlation (|*ρ*| = 0.2). The TOW-CM and MANOVA statistic could benefit from increased traits correlations, and offer much improved power by incorporating strongly correlated null traits. Thus, our results verify the conclusion of Stephens (2013) and Wu and Pankow (2015).

Overall, we can see that the proposed TOW-CM is an attractive approach with good power in most of the scenarios.

## Application to the COPDGene

Chronic obstructive pulmonary disease (COPD) is one of the most common lung diseases characterized by long term poor airflow and is a major public health problem (Murphy and Sethi 2002). The COPDGene Study is a multi-center genetic and epidemiologic investigation to study COPD (Regan et al. 2011). This study is sufficiently large and appropriately designed for analysis of COPD. In this study, we consider more than 5000 non-Hispanic Whites (NHW) participants where the participants have completed a detailed protocol, including questionnaires, pre- and post-bronchodilator spirometry, high-resolution CT scanning of the chest, exercise capacity (assessed by six-minute walk distance), and blood samples for genotyping. The participants were genotyped using the Illumina OmniExpress platform. The genotype data have gone through standard quality-control procedures for genome-wide association analysis detailed at http://www.copdgene.org/sites/default/files/GWAS_QC_Methodology_20121115.pdf.

Based on the literature studies of COPD (Chu et al. 2014 and Han et al. 2011), we selected 7 key quantitative COPD-related phenotypes, including FEV1 (% predicted FEV1), Emphysema (Emph), Emphysema Distribution (EmphDist), Gas Trapping (GasTrap), Airway Wall Area (Pi10), Exacerbation frequency (ExacerFreq), Six-minute walk distance (6MWD), and 4 covariates, including BMI, Age, Pack-Years (PackYear) and Sex. EmphDist is the ratio of emphysema at −950 HU in the upper 1/3 of lung fields compared to the lower 1/3 of lung fields where we did a log transformation on EmphDist in the following analysis, referred to Chu et al. (2014). In the analysis, participants with missing data in any of these phenotypes were excluded.

To evaluate the performance of our proposed method on a real data set, we applied all of the 4 methods (TOW-CM, MANOVA, MSKAT and minP) to the COPD associated genes or genes contain significant single-nucleotide polymorphisms (SNPs) in NHW population with COPD-related phenotypes (Berndt et al. 2012). In the analysis, we first removed the missing data in any genotypic variants and then adjusted each of the 7 phenotypes for the 4 covariates using linear models. In the analysis, participants with missing data in any of the 11 variables were excluded. Therefore, a complete set of 5,430 individuals across 50 genes were used in the following analyses. In order to compare these methods, we adopted the commonly used 10^7^ permutations for TOW-CM and minP methods. For this verification study, we use 0.05 as significance level for MANOVA, MSKAT and TOW-CM method and use Bonferroni corrected significance level 0.05/7 = 7.14 × 10^−3^ for minP methods since this method perform association tests across each trait, respectively. The results are summarized in Table 2. From Table 2, we can see that TOW-CM identified 14 Genes, minP identified 14 Genes, MANOVA identified 12 Genes and MSKAT identified 4 Genes. Among these four methods, TOW-CM identified the most significant genes where all of these 14 genes had previously been reported to be in association with COPD by eligible studies (Berndt et al. 2012; Liang et al. 2016), among which 5 genes (LOC105377462,CHRNA3, CHRNA5, HYKK, IREB2) are statistically significant if we use a more stringent cut-off (**1.00** × **10**^−**3**^) for a multiple testing issue with 50 genes in total.

**Table 2:** The p-values of significant genes in the genetic association analysis for COPD using these four different methods.

| Chr | Genes | Range of MAF | minP<br>(**0.05/7**) | MANOVA<br>(**0.05**) | MSKAT<br>(**0.05**) | TOW-CM<br>(**0.05**) |
| --- | --- | --- | --- | --- | --- | --- |
| 1 | EPHX1 | (0.0214, 0.4620) | 0.0197 | 0.6055 | 0.5890 | 0.6257 |
| 1 | IL6R | (0.1680, 0.4398) | 0.2646 | 0.5214 | 0.5163 | 0.5148 |
| 1 | MFAP2 | (0.1789, 0.4842) | 0.0753 | 0.6986 | 0.9926 | 0.6869 |
| 1 | TGFB2 | (0.0139, 0.4858) | **$7.23 \times 10^{-4}$** | 0.2282 | 0.1831 | **$3.47 \times 10^{-2}$** |
| 2 | HDAC4 | (0.0147, 0.4906) | 0.0468 | 0.3393 | 0.2197 | 0.5026 |
| 2 | SERPINE2 | (0.0143, 0.4642) | 0.4671 | 0.9797 | 0.7706 | 0.9010 |
| 2 | SFTPB | (0.0784, 0.4766) | 0.0738 | 0.1017 | 0.1669 | 0.3921 |
| 2 | TNS1 | (0.0128, 0.4936) | 0.00727 | **$4.63 \times 10^{-2}$** | 0.2095 | **$2.65 \times 10^{-2}$** |
| 3 | MECOM | (0.0099, 0.4957) | 0.0359 | 0.9878 | 0.7211 | 0.9735 |
| 3 | RARB | (0.0278, 0.4942) | 0.0491 | 0.1988 | 0.7469 | 0.3973 |
| 4 | LOC105377462 | (0.0190, 0.4933) | **0.00** | **$6.28 \times 10^{-3}$** | 0.8310 | **0.00** |
| 4 | FAM13A | (0.0279, 0.4968) | **$1.08 \times 10^{-5}$** | 0.2169 | 0.0939 | 0.3925 |
| 4 | GC | (0.0511, 0.4397) | 0.1743 | 0.1875 | 0.6499 | 0.5257 |
| 4 | GSTCD | (0.0343, 0.3872) | **$3.6 \times 10^{-6}$** | **$5.63 \times 10^{-5}$** | 0.1376 | **$3.30 \times 10^{-2}$** |
| 4 | HHIP | (0.0368, 0.4984) | 0.0150 | **$3.64 \times 10^{-2}$** | 0.4131 | **$1.95 \times 10^{-3}$** |
| 5 | HTR4 | (0.0396, 0.4889) | 0.0487 | 0.6622 | 0.6512 | 0.8906 |
| 5 | SPATA9 | (0.1059, 0.4077) | 0.1145 | 0.3118 | 0.5198 | 0.1964 |
| 6 | TNF | (0.0259, 0.0809) | 0.0320 | 0.1627 | 0.1542 | 0.3077 |
| 6 | ZKSCAN3 | (0.0137, 0.3036) | 0.4990 | 0.8575 | 0.9083 | 0.8793 |
| 6 | AGER | (0.0442, 0.1830) | **$3.25 \times 10^{-4}$** | **$2.27 \times 10^{-3}$** | **$9.31 \times 10^{-4}$** | **$9.57 \times 10^{-3}$** |
| 6 | ARMC2 | (0.0187, 0.4695) | 0.1618 | 0.2481 | 0.1474 | 0.6233 |
| 6 | NCR3 | (0.0133, 0.0899) | 0.0735 | 0.4641 | 0.4145 | 0.5892 |
| 6 | SOX5 | (0.0193, 0.4972) | 0.0764 | 0.8376 | 0.6845 | 0.6386 |
| 10 | LRMDA | (0.0094, 0.4956) | 0.0394 | 0.4102 | 0.7190 | 0.3260 |
| 10 | CDC123 | (0.0240, 0.4561) | 0.0138 | 0.6846 | 0.4097 | 0.8836 |
| 10 | GSTO2 | (0.0538, 0.4547) | **$4.0 \times 10^{-7}$** | **$1.36 \times 10^{-6}$** | 0.1387 | 0.8731 |
| 10 | SFTPD | (0.0186, 0.4367) | 0.3699 | 0.9997 | 0.9767 | 0.9751 |
| 11 | GSTP1 | (0.3351, 0.3452) | **$6.60 \times 10^{-3}$** | 0.7053 | 0.1211 | 0.5043 |
| 11 | MMP1 | (0.0519, 0.3916) | 0.1665 | 0.8614 | 0.6557 | 0.9449 |
| 11 | MMP12 | (0.0541, 0.1439) | 0.4073 | 0.9512 | 0.7372 | 0.8941 |

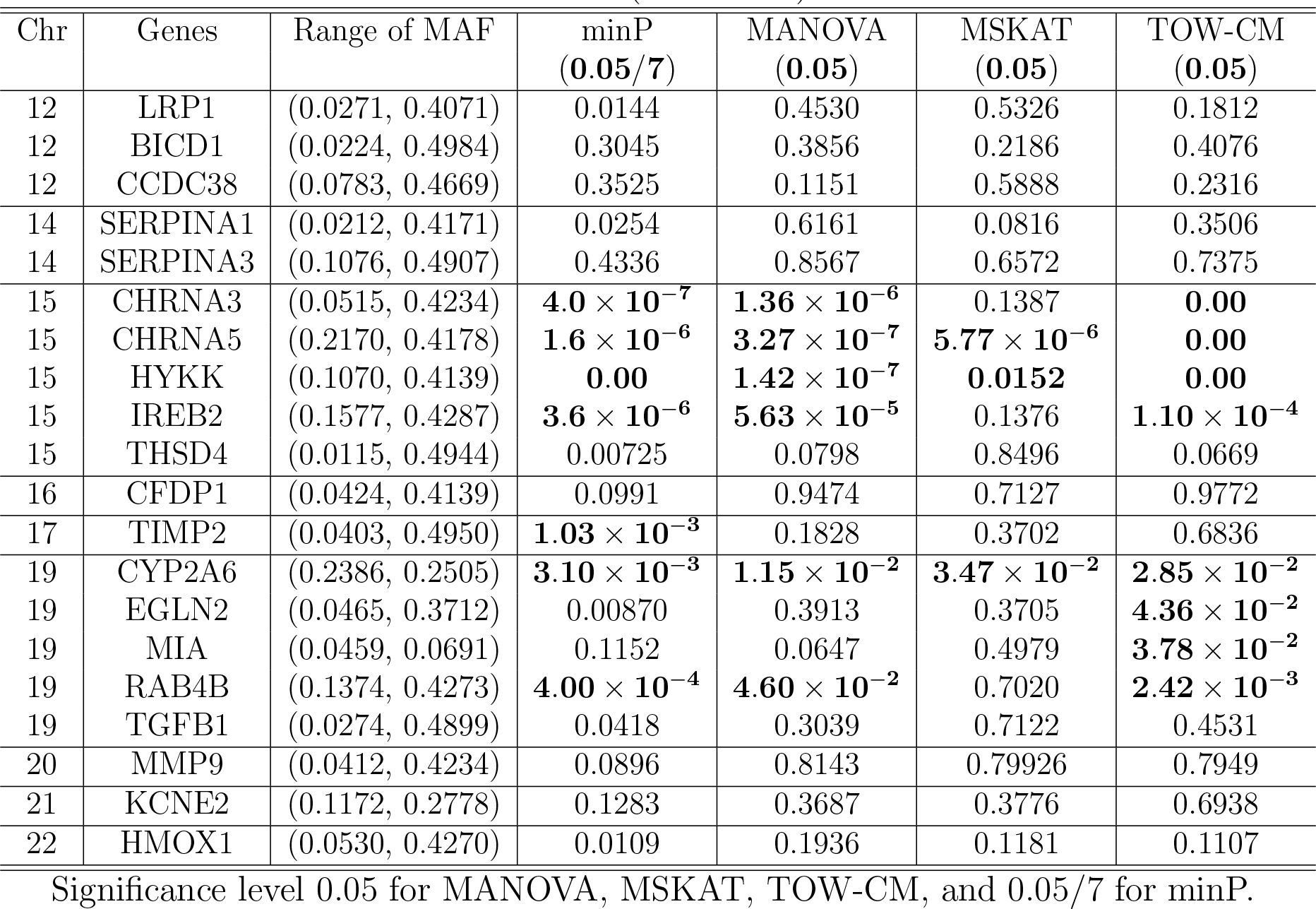

## Discussion

GWAS have identified many variants with each variant affecting multiple phenotypes, which suggests that pleiotropic effects on human complex phenotypes may be widespread. Also, recent studies have shown that complex diseases are caused by both common and rare variants (Bodmer and Bonilla 2008; Pritchard and Cox 2002; Teer and Mullikin 2010). Therefore, statistical methods that can jointly analyze multiple phenotypes for common or/and rare variants have advantages over analyzing each phenotype individually or only considering for common variants (GWAS). In this article, we propose TOW-CM method to perform multivariate analysis for multiple phenotypes in association studies based on the following reasons: (1) complex diseases are usually measured by multiple correlated phenotypes in genetic association studies; there is increasing evidence showing that studying multiple correlated phenotypes jointly may increase power for detecting disease associated genetic variants, and (3) there is a shortage of gene-based approaches for multiple traits. Simulation results show that TOW-CM has correct type I error rates and is consistently more powerful than the other four compared tests. The real data analysis results show that TOW-CM has excellent performance in identifying genes associated with complex disease with multiple correlated phenotypes such as COPD.

One disadvantage of TOW-CM is that the test statistic does not have an asymptotic distribution and a permutation procedure is needed to estimate its P-value, which is time consuming compared to the methods whose test statistics have asymptotic distributions. To save time when applying TOW-CM to genetic association studies, we can use the step-up procedure (Pan et al. 2014) to determine the number of permutations, which can show evidence of association based on a small number of permutations first (e.g. 1,000) and then a large number of permutations are used to test the selected potentially significant genes. Furthermore, TOW-CM method can not only be used for gene-based association studies, but also can be extended to pathway-based analysis or even transcriptome-wide association study (TWAS). How to extend our method to pathway-based analysis or TWAS is our future work.

## Acknowledgments

Q Sha was supported by the National Human Genome Research Institute of the National Institutes of Health under Award Number R15HG008209. The content is solely the responsibility of the authors and does not necessarily represent the official views of the National Institutes of Health.

The Genetic Analysis workshops are supported by NIH grant R01 GM031575 from the National Institute of General Medical Sciences. Preparation of the Genetic Analysis Workshop 17 Simulated Exome Data Set was supported in part by NIH R01 MH059490 and used sequencing data from the 1000 Genomes Project (www.1000genomes.org).

This research used data generated by the COPDGene study (phs000179/HMB and phs000179 /DS-CS-RD), which was supported by National Institutes of Health (NIH) grants U01HL089856 and U01HL089897. The content is solely the responsibility of the authors and does not necessarily represent the official views of the National Heart, Lung, and Blood Institute or the National Institutes of Health. The COPDGene project is also supported by the COPD Foundation through contributions made by an Industry Advisory Board comprised of Pfizer, AstraZeneca, Boehringer Ingelheim, Novartis, and Sunovion.

